# Test-retest reliability and long-term stability of 3-tissue constrained spherical deconvolution methods for analyzing diffusion MRI data

**DOI:** 10.1101/764506

**Authors:** Benjamin T. Newman, Thijs Dhollander, Kristen A. Reynier, Matthew B. Panzer, T. Jason Druzgal

## Abstract

**Purpose:** Several recent studies have utilized a 3-tissue constrained spherical deconvolution pipeline to obtain quantitative metrics of brain tissue microstructure from diffusion-weighted MRI data. The three tissue compartments, comprising white matter-, grey matter-, and CSF-like (free water) signals, are potentially useful in the evaluation of brain microstructure in a range of pathologies. However, the reliability and long-term stability of these metrics has not yet been evaluated.

**Methods:** This study examined estimates of whole brain microstructure for the three tissue compartments, in three separate test-retest cohorts. Each cohort has different lengths of time between baseline and retest, ranging from within the same scanning session in the shortest interval to three months in the longest interval. Each cohort was also collected with different acquisition parameters.

**Results:** The CSF-like compartment displayed the greatest reliability across all cohorts, with intraclass correlation coefficient (ICC) values being above 0.95 in each cohort. White matter-like and grey matter-like compartments both demonstrated very high reliability in the immediate cohort (both ICC>0.90), however this declined in the 3 month interval cohort to both compartments having ICC>0.80. Regional CSF-like signal fraction was examined in bilateral hippocampus and had an ICC>0.80 in each cohort.

**Conclusion:** The 3-tissue CSD techniques provide reliable and stable estimates of tissue microstructure composition, up to 3 months longitudinally in a control population. This forms an important basis for further investigations utilizing 3-tissue CSD techniques to track changes in microstructure across a variety of brain pathologies.

## Introduction

Diffusion-weighted Magnetic Resonance Imaging (dMRI) is a widely used, noninvasive, method for measuring the diffusion of water molecules in the brain. Within the microarchitectural environment of the brain, diffusion of water molecules is hindered by various cellular components, particularly the lipid bilayers that make up cell membranes. This principle has been applied to study white matter fiber bundles (“tracts”), as the myelin sheaths surrounding neuronal axons result in anisotropic diffusion (Basser et al., 1994; Pierpaoli & Basser, 1996; Boullerne, 2016). dMRI has seen widespread use in studies of brain connectivity as well as in clinical populations and neurosurgery (Le Bihan et al., 2001; Nimsky et al., 2005; Alexander et al., 2007; Johansen-Berg & Behrens, 2013).

Initially, anisotropic diffusion was typically modelled using a tensor, which sought to quantify both the average orientation, anisotropy, and magnitude of diffusion within each voxel of the brain; this approach is known as Diffusion Tensor Imaging (DTI, Basser et al., 1994). More recently, the dMRI modelling domain has seen a proliferation in novel, more advanced, mathematical methods for analyzing the diffusion-weighted signal. These methods aim to overcome several shortcomings of applying the relatively simplistic DTI model to the complex diffusion-weighted signals observed in the brain. This complexity primarily arises from two physiological qualities of the brain itself: the first being crossing fibers, where white matter (WM) tracts occupying the same voxel are oriented differently in space (Wiegell et al., 2000; Tuch et al., 2003); and the second being the presence of other fluids and tissues, including cerebrospinal fluid (CSF) and grey matter (GM) and other cell bodies which “contaminate” the directional signal (Chenevert et al., 1990; Jones & Cercignani, 2010; Rydhög et al, 2017). These are major issues as it has been estimated that up to 90% of WM tissue voxels contain more than one WM fiber tract orientation (Jeurissen et al., 2013), and partial voluming effects alone ensure that a substantial number of voxels contain proportions of multiple tissue and/or fluid compartments (Alexander et al., 2001).

To address these issues, and with the advent of high angular resolution diffusion imaging (HARDI) acquisition protocols, more advanced methods for describing the observed dMRI data have been proposed by a number of researchers (Tournier et al., 2011; for reviews see Assaf & Pasternak, 2008; and Dell’Acqua & Tournier 2018). One such method, Constrained Spherical Deconvolution (CSD), allows for the presence of multiple fibers along different orientations (Tournier et al., 2007). CSD resolves these orientations by deconvolving the signal profile corresponding to a prototypical single fiber-like voxel (termed a response function) from the observed signal in each and every other voxel, resulting in the orientation of fibers as a continuous angular function termed the Fiber Orientation Distribution (FOD). Quantitative information can also be obtained from the FOD, as a measure of “Apparent Fiber Density” (AFD) for each fiber population (Raffelt et al., 2012).

The original (“single-tissue”) CSD has been expanded into Multi-Shell Multi-Tissue CSD (MSMT-CSD) by performing a similar deconvolution with 3 separate WM, GM, and CSF-like tissue response functions. The approach was initially aimed at separating signal originating from GM and CSF-like tissue compartments, in order to improve the accuracy of the WM FOD itself, which otherwise appears very noisy (with many false positive “peaks” or lobes) when using single-tissue CSD in areas of partial voluming with other tissues and fluids (Jeurissen et al., 2014). This subsequently benefits several other analysis and processing steps, such as streamline tractography, which heavily rely on a “clean” and accurate WM FOD. MSMT-CSD thus attempted to address the main shortcomings of the DTI model as well as additional remaining shortcomings of single-tissue CSD.

As its name hints at, MSMT-CSD requires a *multi-shell* diffusion acquisition scheme in order to successfully tease apart contributions from the 3 WM-, GM- and CSF-like compartments at once. However, to obtain the same benefits offered by MSMT-CSD, yet using only *single-shell* data, Dhollander & Connelly, (2016) have proposed a novel approach named Single-Shell 3-Tissue CSD (SS3T-CSD) that can resolve the WM-, GM- and CSF-like compartments as well. By relying only on *single-shell* data, it allows for shorter acquisition times and is compatible with a wider range of data, both historical as well as clinical.

Resolving these different compartments using either 3-tissue CSD method (i.e., MSMT-CSD or SS3T-CSD) holds value beyond improving WM tractography: it can also serve as a proxy for the evaluation of brain microstructure and tissue composition (Dhollander et al., 2017; Mito et al., 2018; Mito et al. 2019). By interrogating brain voxels for diffusion signal patterns that look ‘like’ compositions of the diffusion signals represented by the WM/GM/CSF response functions, it might be possible to gain quantifiable information about microstructure (Figure 1). Using these basic compartments as a diffusion signal model focuses more on coarse properties of brain tissue microstructure rather than separating similar cell types (e.g. different populations of glial cells), or separating different types of pathology (e.g. edema, CSF-infiltration in neurodegeneration, and damage from ischemic stroke). Although, provided with a known context, reasonable inferences of such pathology might be possible to make nonetheless. Even for WM tractography in cases of infiltration by pathological tissues, the 3-tissue CSD approach can provide direct benefits in terms of recovering healthy WM structures, e.g. in infiltrating tumors (Aerts et al. 2019).

**Figure 1:**
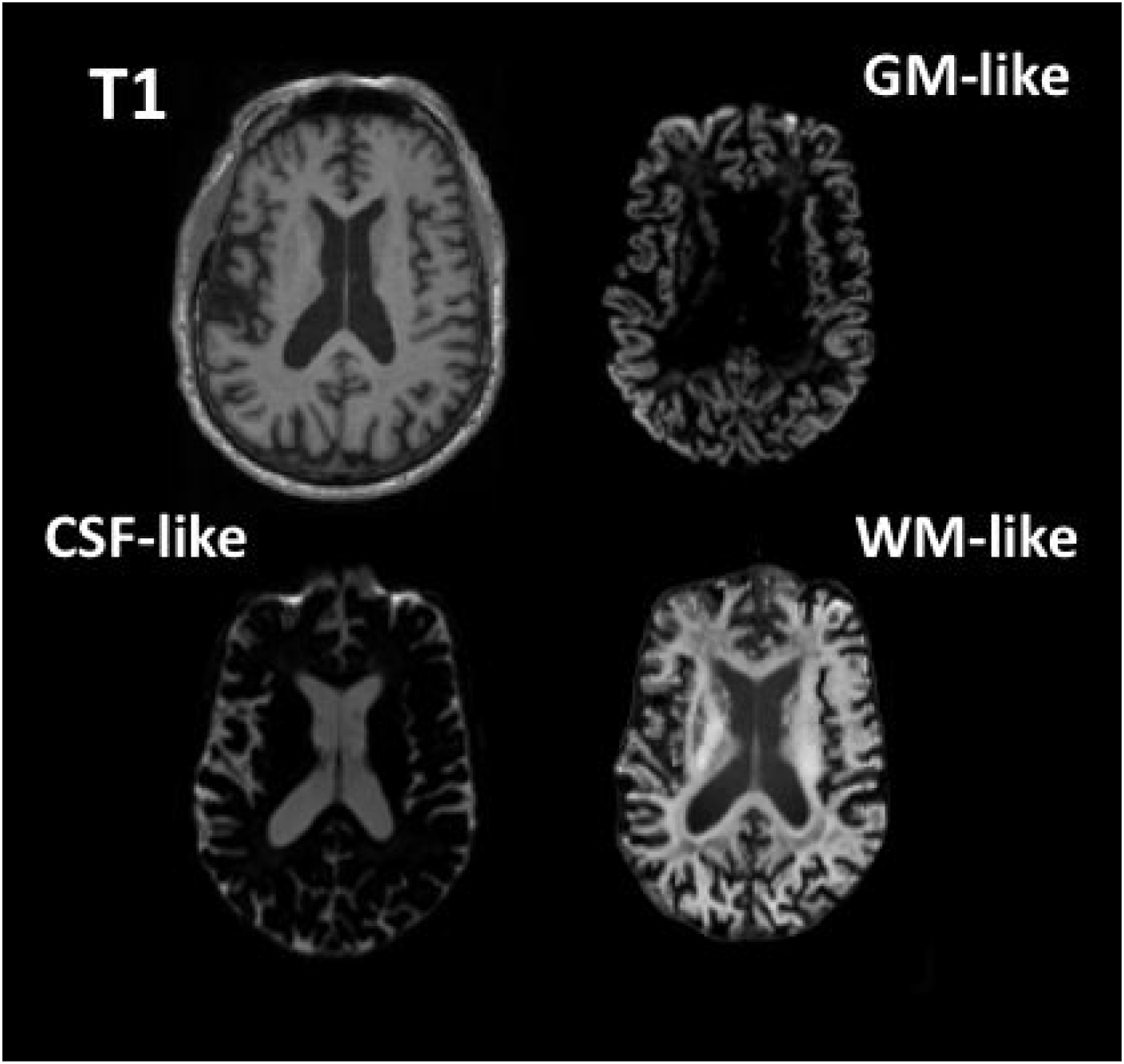
Axial slices showing a T1-weighted MPRAGE and the GM-, CSF-, and WM-like tissue compartments derived from the dMRI data using 3-tissue CSD.

3-tissue CSD derived compartments are a promising, non-invasive method for exploring tissue composition in the brain. The utilization of this approach toward analyzing tissue composition might hold advantages over tensor-based models such as Free Water Elimination (FWE, Pasternak et al., 2009). The free water estimate from the FWE technique was shown to have limited reproducibility: errors ranged from 5.2-18.2% across ROIs in a test-retest cohort (Albi et al., 2017). The CSF-like compartment from 3-tissue CSD techniques might provide an alternative way to recover free water contribution to the signal, using a WM model that does take into account crossing fibres (as opposed to a tensor method). With the advances provided in SS3T-CSD, it is also able to provide signal contribution from the full 3 tissue compartments using *single-shell* data (i.e. equivalent to acquisition requirements for the FWE technique), allowing for a broader range of input data compared to other 3-tissue compartment models such as NODDI (Zhang et al., 2012). In a recent review of microstructural diffusion imaging applied to psychiatric disorders, Pasternak et al., (2018) illustrated the acquisition sequence complexity compared to the number of microstructure compartments evaluated for several common dMRI analysis techniques. Addition of MSMT-CSD and SS3T-CSD illustrate the range of data required for input to a range of models and the capabilities of resolving compartments compared to other techniques (Figure 2).

**Figure 2:**
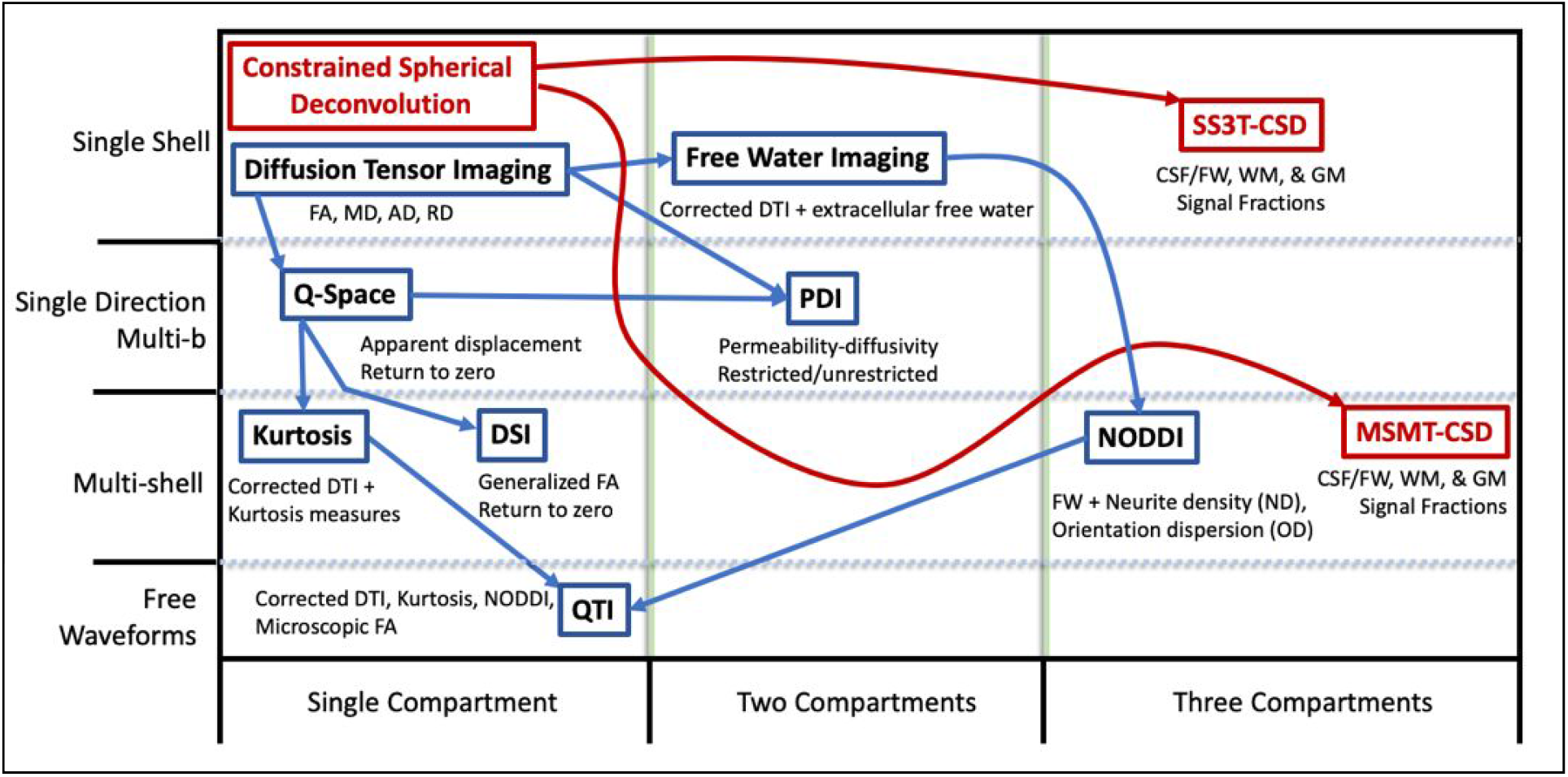
Chart adapted from Pasternak et al., (2018); comparison of common DTI and other model metrics to CSD derived tissue signal fractions by requirements of acquisition (rows) and number of output compartments (columns). Methods derived from CSD have been added in red.

To date, there has not been a quantitative test-retest study examining the reliability and long-term stability of 3-tissue CSD techniques. The purpose of this study is to provide evidence that 3-tissue CSD techniques are a reliable and stable approach for assessing brain microarchitecture, via analysis of the 3 resulting tissue signal fractions.

## Methods

### Cohorts

Three test-retest cohorts were retrospectively evaluated in this study: two local datasets collected at the University of Virginia from ongoing research projects, and one publicly available dataset obtained from the Nathanial Kline Institute for Psychiatric Research: enhanced test-retest (eNKI-TRT) as part of the 1000 Functional Connectomes Project (Biswal et al., 2010; Nooner et al., 2012). Both studies collected at the University of Virginia received ethical approval from the University of Virginia Institutional Review Board for Health Sciences Research. Each cohort has different time intervals between baseline and retest scans, and was collected with different acquisition parameters. This approach allows reliability to be measured under conditions that represent a variety of different diffusion imaging parameters. Examining stability across different time periods allows for insight into the potential for longitudinal studies tracking changes in 3-tissue signal fractions in individuals or between groups over time.

The first cohort (”*immediate rescan*” cohort) examined immediate test-retest reliability by performing identical dMRI acquisitions sequentially without table repositioning. This cohort consisted of individuals participating in a separate study at the University of Virginia that included multiple scanning sessions. The cohort consisted of 20 healthy control participants (all male, age at baseline: 22.8±3.0 SD). Each participant was scanned twice at each of 3 visits (with the exception of one participant who only attended 2 scans) for a total of 59 baseline-rescan pairs collected for analysis.

The second cohort (”*short timescale*” cohort) is representative of the quality of diffusion imaging found in large-scale, open science cohorts. Subjects were selected from the original NKI Rockland community study, a group intentionally recruited for similarity to the demographics of the broader United States as a whole (Nooner et al., 2012). 20 subjects (5 female, age at baseline: 34.4±12.9 SD) had diffusion MRI data available at both baseline and rescan. All participants were rescanned within a range of 7-60 days after baseline. Subjects were *not* excluded for any history of illness, and 2 participants had a diagnosed history of prior alcohol abuse while 2 other participants had a diagnosed history of a major depressive disorder. Both of these diagnoses are known to affect brain function and structure (Oscar-Berman & Marinkovic, 2007; Jiang et al., 2017) but the nature of the within-subjects design did not necessitate removing any individuals from the study.

The third cohort (”*long timescale*” cohort) was collected as a healthy control group for a previously published study conducted at the University of Virginia examining college athletes (Reynolds et al., 2017). 52 participants (all male, age at baseline: 21.9±3.3 SD) were re-scanned 3-4 months after baseline (mean days between scans: 107.9±7.1 SD) and were screened for a history of neurologic disease or concussion.

### Image Acquisition

As discussed previously, data from the three cohorts were acquired using different protocols.

The *immediate rescan* cohort was scanned using a Siemens Prisma 3T scanner with an isotropic voxel size of 1.7×1.7×1.7mm^3^, TE=70ms and TR=2900ms. Using a multi-shell protocol, 10 b=0 images and 64 gradient directions at both b=1500s/mm^2^ and b=3000s/mm^2^ were acquired. This protocol was applied twice with one immediately following the other without actively repositioning the participant in the scanner.

The *short timescale* cohort was acquired externally and obtained through the Neuroimaging Tools and Resources Collaboratory at www.nitrc.org. Imaging data was collected using a Siemens Trio Tim with an isotropic voxel size of 2×2×2mm^3^, TE=85ms and TR=2400ms. Using a single-shell protocol, 9 b=0 images and 127 gradient directions at b=1500s/mm^2^ were acquired.

The *long timescale* cohort was scanned using the same Siemens Prisma 3T scanner as the first (immediate rescan) cohort using a different protocol with an isotropic voxel size of 2.7×2.7×2.7mm^3^, TE=100ms. Using a multi-shell protocol, 1 b=0 image and 30 gradient directions at both b=1000s/mm^2^ and b=2000s/mm^2^ were acquired.

### Analysis

Data preprocessing was largely identical across all images in all cohorts in the study. Images were first denoised (Veraart et al., 2016). Gibbs ringing was then corrected (Kellner et al., 2016). This was followed by utilizing the FSL package (*“topup”* and *“eddy”*) to correct for susceptibility induced (EPI) distortions, eddy currents, and subject motion (Smith et al., 2004; Andersson et al., 2003; Andersson & Sotiropoulos, 2016; Andersson et al., 2016). Finally, we upsampled the preprocessed data to 1.3×1.3×1.3mm^3^ isotropic voxels (Greenspan, 2008; Kuklisova-Murgasova et al., 2012; Bastiani et al., 2019). These preprocessing steps are largely similar to those used in other recently published works (Bastiani et al., 2019; Pietsch et al., 2019; Mito et al., 2019; Aerts et al., 2019). Brain masks were obtained for all subjects by performing a recursive application of the Brain Extraction Tool (Avants et al., 2014).

For 3-tissue CSD processing, the 3-tissue response functions were obtained from the data themselves using an unsupervised method (Dhollander et al., 2016), resulting in the single-fiber WM response function as well as isotropic GM and CSF response functions for each subject. For each tissue type (WM, GM, CSF), the response function was averaged across all individuals in each cohort to obtain a single unique set of 3-tissue response functions per cohort. For the *multi-shell* data in the immediate rescan and long timescale cohorts, MSMT-CSD was performed (Jeurissen et al., 2014). For the *single-shell* data in the short timescale cohort, SS3T-CSD was performed (Dhollander & Connelly, 2016). For all subjects in all cohorts, this resulted in their WM-like compartment (represented by a complete WM FOD) as well as GM-like and CSF-like compartments. The CSF-like compartment can in this context also be interpreted as a free-water (FW) compartment (Dhollander et al., 2017). Finally, each subject’s three tissue compartments were then normalised to sum to 1 on a voxel-wise basis, resulting in the final 3-tissue signal *fraction* maps (Mito et al., 2019); the metrics for which we performed the test-retest analyses in this work.

To measure the CSF-like (free water) signal fraction in the hippocampus of each subject in the immediate rescan cohort, a template was first produced from a combination of all 118 individual scans. This was achieved using an affine, followed by non-linear registration on the WM FODs themselves in an unbiased manner (Raffelt et al., 2011). A whole brain WM image from the LONI atlas (Shattuck et al., 2008) was registered along with each hippocampus map to the template using the ANTs image registration toolbox (Avants et al., 2009) and then subsequently warped to each individual scan using the reverse transform from template creation. In native space, an average was computed of the CSF-like (free water) signal fractions in the ROI, using only voxels with a CSF-like signal fraction smaller than 0.5, to mimic free water analysis (i.e., to avoid accidentally including voxels outside of the brain parenchyma, which might be entirely CSF-filled spaces).

All processing was performed using a combination of different software packages: MRtrix3 (Tournier et al., 2019), MRtrix3Tissue (https://3Tissue.github.io, a fork of MRtrix3), FSL (Jenkinson et al., 2012), ANTs (Avants et al., 2009).

## Results

The CSF-like (free water) tissue signal fraction map was restricted to voxels where the corresponding WM and GM signal maps summed to greater than 50%. This allowed for analysis of the CSF-like signal fraction in tissue without including the ventricles or subarachnoid space, the bulk size of which would otherwise bias a proper whole-brain free water measurement. Additionally, the CSF-like infiltration into brain tissue is a potentially more interesting measurement in the context of healthy functioning or pathology; and is indeed designed to be comparable to measurements of free water encountered in the literature (Pasternak et al., 2009). For all cohorts, results from the 3-tissue signal fractions were averaged across the brain parenchyma. Averages for baseline and retest values were compared by calculating the intraclass correlation coefficient (ICC) and Pearson’s correlations. The results for both of these measures are summarized in Table 1.

**Table 1:** Statistical analysis of the 3 test-retest cohorts in the experiment; p-values are calculated based on the Pearson’s correlation. For each cohort the left hippocampus (LH) and right hippocampus (RH) were selected as ROIs and the CSF-like (free water) signal fraction was measured to examine the reliability of 3-tissue CSD derived free water estimates in subcortical structures specifically as well.

| Dataset | Tissue | Subjects | ICC | Pearson's Rho | p-value |
| --- | --- | --- | --- | --- | --- |
| *Immediate rescan* | CSF-like | 59 | 0.9731 | 0.9636 | <0.001 |
|  | WM-like | 59 | 0.9868 | 0.9748 | <0.001 |
|  | GM-like | 59 | 0.9929 | 0.9868 | <0.001 |
|  | LH CSF-like | 59 | 0.9578 | 0.9181 | <0.001 |
|  | RH CSF-like | 59 | 0.9376 | 0.8915 | <0.001 |
| *Short timescale (7-60 days)* | CSF-like | 20 | 0.9546 | 0.9281 | <0.001 |
|  | WM-like | 20 | 0.9692 | 0.9423 | <0.001 |
|  | GM-like | 20 | 0.9852 | 0.9700 | <0.001 |
|  | LH CSF-like | 20 | 0.9332 | 0.9169 | <0.001 |
|  | RH CSF-like | 20 | 0.9094 | 0.8469 | <0.001 |
| *Long timescale (3-4 months)* | CSF-like | 52 | 0.9564 | 0.9364 | <0.001 |
|  | WM-like | 52 | 0.8157 | 0.7200 | <0.001 |
|  | GM-like | 52 | 0.8746 | 0.8024 | <0.001 |
|  | LH CSF-like | 52 | 0.8516 | 0.7421 | <0.001 |
|  | RH CSF-like | 52 | 0.8217 | 0.7118 | <0.001 |

Specific test-retest correlations for each of the three tissue types derived from the 3-tissue CSD techniques are presented in Figures 3–5. All correlations between baseline and retest were significant in all cohorts; the highest whole brain ICC values were obtained from the *immediate rescan* cohort (Figure 3). In the *short timescale* cohort, similar to the immediate rescan cohort, all compartments had an ICC value above 0.95 and Pearson’s Rho above 0.90 (Figure 4). The *long timescale* cohort had slightly declined performance, yet with the ICC value for all compartments still being larger than 0.80 (Figure 5).

**Figure 3:**
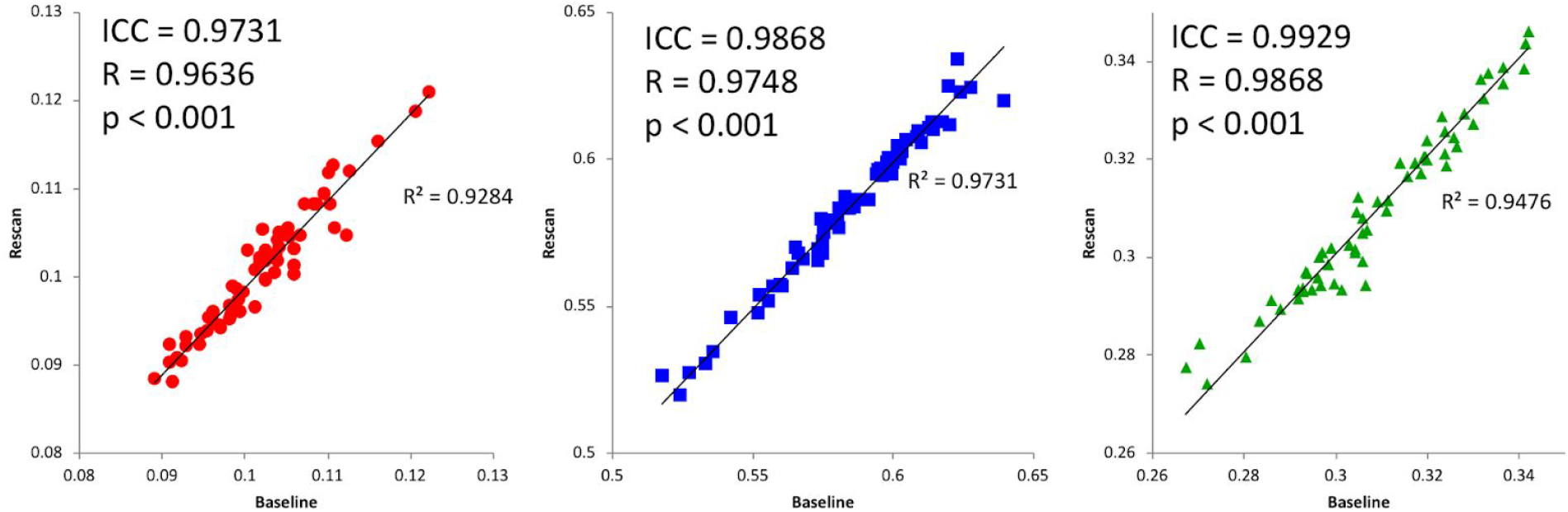
*Immediate rescan* baseline and re-scan values for CSF- (left), WM- (center), and GM-like (right) signal fractions obtained from a cohort scanned with a duplicate sequence immediately following baseline. Includes ICC and Pearson’s correlation values.

**Figure 4:**
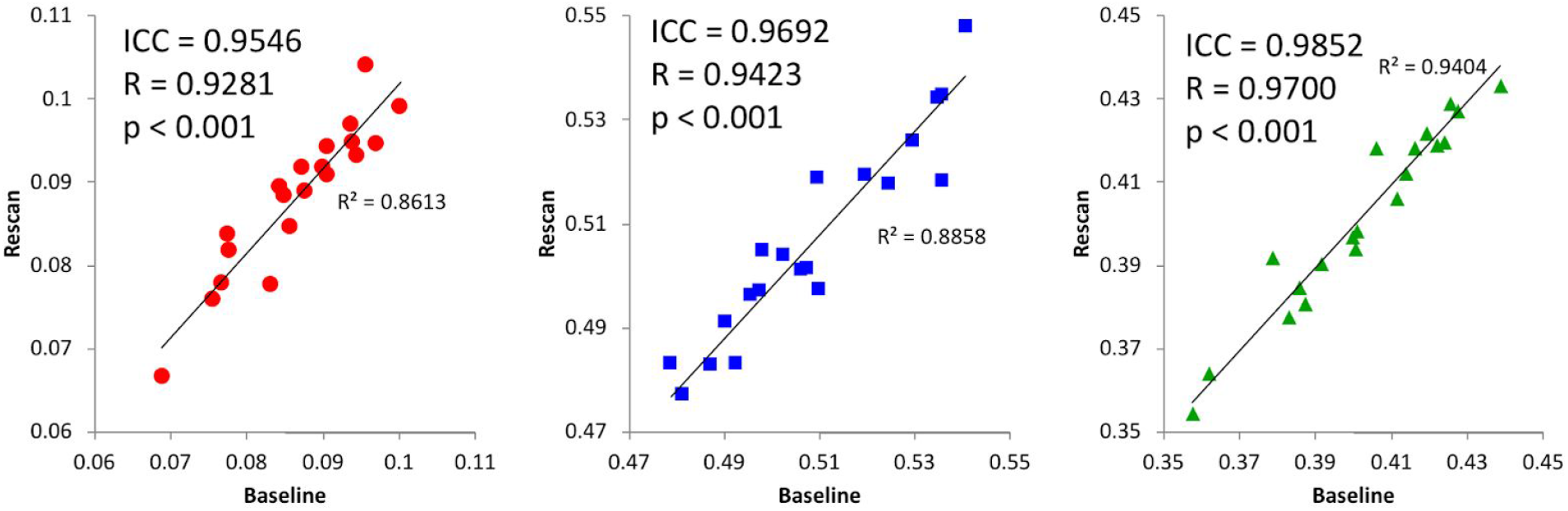
*Short timescale* baseline and re-scan values from CSF- (left), WM- (center), and GM-like (right) signal fractions obtained from a cohort with 7-60 days between baseline and re-scan. Subjects were taken from the eNKI group and their single-shell dMRI data analyzed with SS3T-CSD. Includes ICC and Pearson’s correlation values.

**Figure 5:**
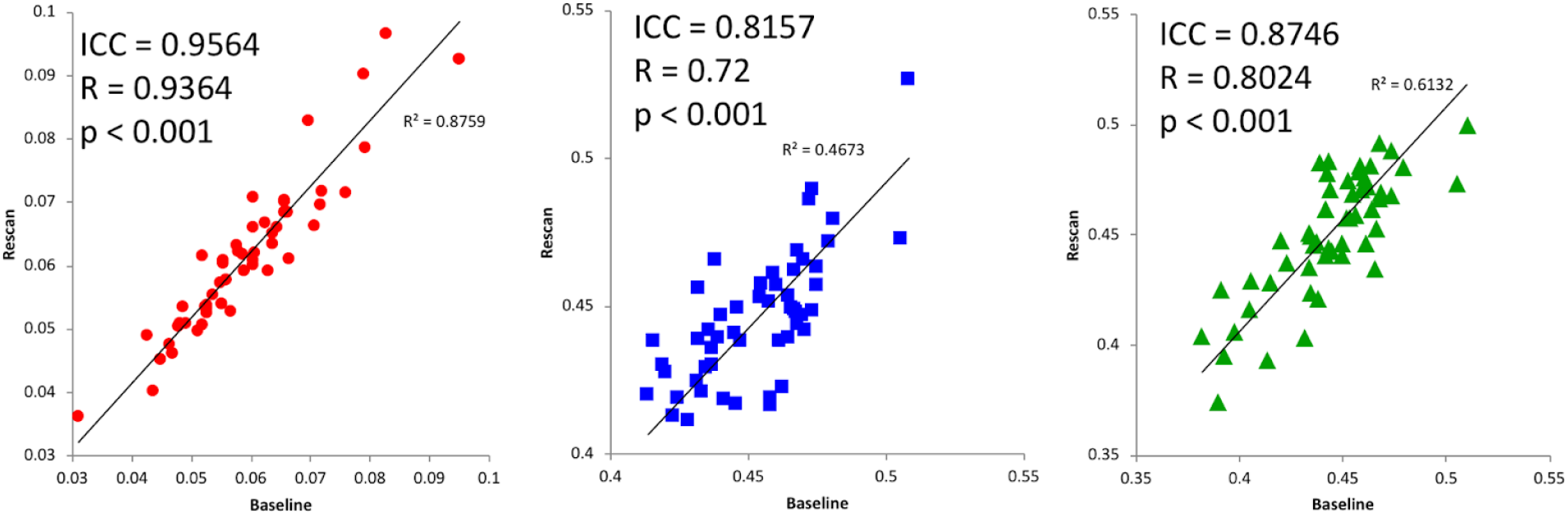
*Long timescale* baseline and re-scan values from CSF- (left), WM- (center), and GM-like (right) signal fractions obtained from a cohort with 3 months between baseline and re-scan. Includes ICC and Pearson’s correlation values.

In each cohort, the hippocampi were also analyzed separately in order to demonstrate the utility of a 3-tissue CSD approach in a specific region of interest. Bilateral hippocampus was selected for this demonstration as a commonly studied brain ROI with representation from each of the three tissue compartments examined. Comparison of average CSF-like (free water) signal fraction in this ROI between baseline and retest in the *immediate rescan* cohort resulted in an ICC value above 0.90 in both left and right hippocampus, as well as a significant Pearson’s correlation (Figure 6). In the *short timescale* cohort both left and right hippocampus similarly had an ICC value above 0.90 and a significant Pearson’s correlation (Figure 7). In the *long timescale* cohort the reliability was reduced with an ICC value above 0.80 but a still significant Pearson’s correlation (Figure 8).

**Figure 6:**
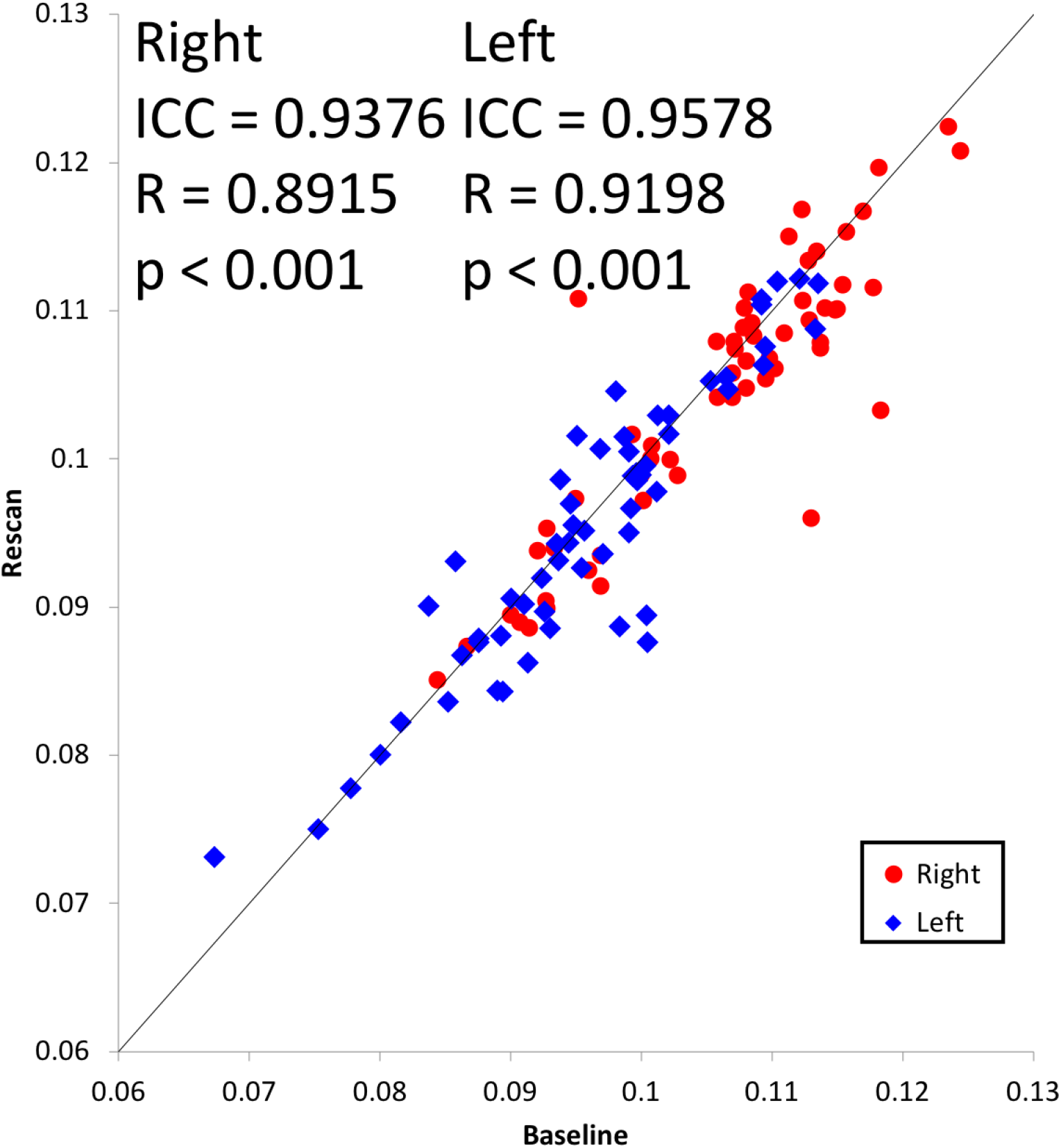
CSF-like signal fraction for the left and right hippocampus in the 59 pairs of baseline-retest scans in the *immediate rescan* cohort. Values for the right hippocampus of each individual are shown in red and values for the left hippocampus are shown in blue.

**Figure 7:**
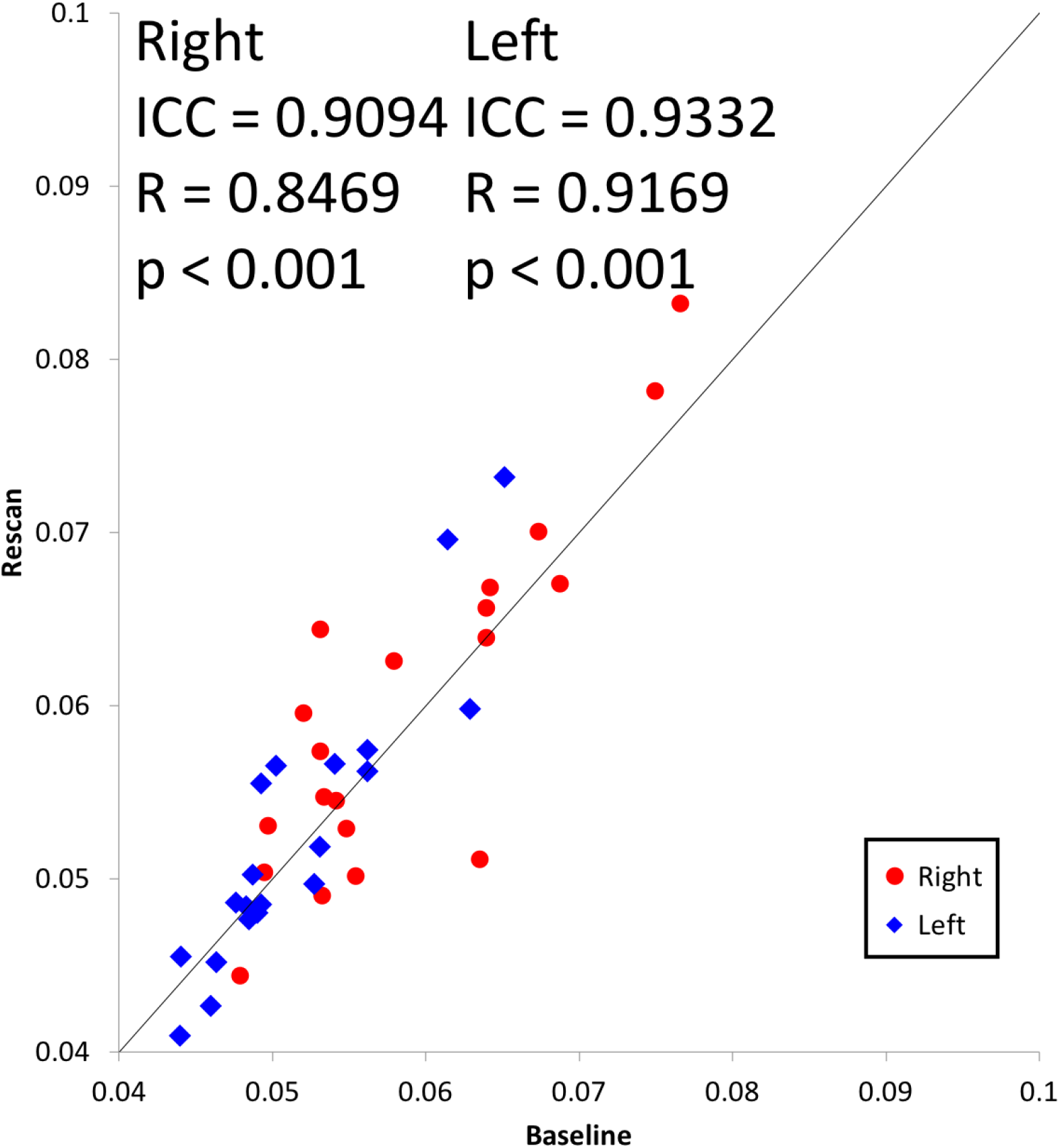
CSF-like signal fraction for the left and right hippocampus in the 20 pairs of baseline-retest scans in the *short timescale* cohort. Values for the right hippocampus of each individual are shown in red and values for the left hippocampus are shown in blue.

**Figure 8:**
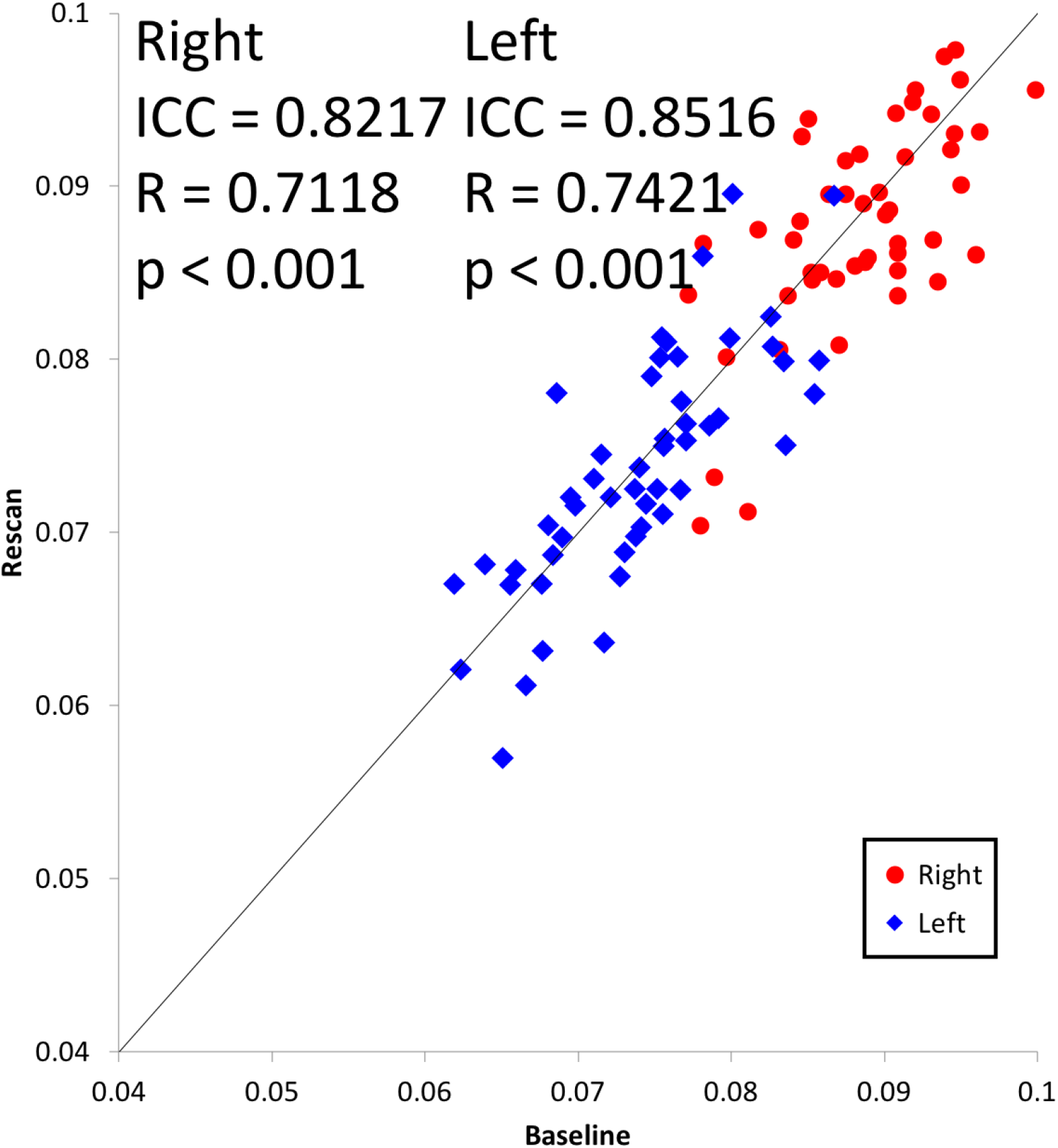
CSF-like signal fraction for the left and right hippocampus in the 52 pairs of baseline-retest scans in the *long timescale* cohort. Values for the right hippocampus of each individual are shown in red and values for the left hippocampus are shown in blue.

There was a consistent asymmetrical effect observed between the CSF-like signal fraction in right and left hippocampus across all cohorts. The CSF-like signal fraction in each subject’s right and left hippocampus were averaged between baseline and rescan and a paired t-test performed for each cohort. This showed that there was a significantly greater CSF-like signal fraction in the right versus the left hippocampus (T_58_ = −10.022, p<0.001; T_19_ = −6.002, p<0.001; and T_51_ = −23.486, p<0.001; for the *immediate rescan, short timescale*, and *long timescale* cohorts, respectively).

## Discussion

Each of the 3-tissue signal fractions demonstrated good reliability across all of the measured timescales we assessed in this work. ICC values were above 0.95 for each of the tissue compartments included in the *immediate rescan* and *short timescale* cohorts. This occurred despite the *short timescale* cohort being *single-shell* data with a b-value of 1500s/mm^2^, and a lower voxel size compared to the other two cohorts (both of which were *multi-shell* and had highest b-values of b=3000s/mm^2^ and b=2000s/mm^2^ for the *immediate timescale* and *long timescale* cohorts, respectively). This result suggests that 3-tissue CSD techniques can reliably obtain quantitative measurements across a range of diffusion imaging protocols, including from openly available datasets. This performance, however, declined slightly in the *long timescale* cohort: the CSF-like (free water) signal fraction within tissue still had an ICC value above 0.95 while the WM-like and GM-like signal fractions had a slightly lower ICC value, yet still above 0.80 nonetheless. Regardless, all Pearson’s correlations were highly significant, indicating that 3-tissue CSD techniques are still able to obtain reliable measurements of brain tissue microstructure, stable up to 3 months from baseline.

Additionally, the free water signal fraction demonstrated good reliability in both hippocampi at each of the examined timescales. ICC values were above 0.80 and a significant effect of laterality was observed consistently across each cohort, with the right hippocampus having significantly higher free water signal fraction than the left hippocampus. Though this study does not suggest any hypothesis for why this laterality was observed, it is consistent with volumetric MRI findings that demonstrate hippocampal asymmetry (for a meta-analysis see Pedraza et al., 2004), as well as a recent study that reported asymmetry in hippocampal free water content (Ofori et al., 2019). This suggests that free water signal fraction is both a reliable quantitative measurement for subcortical ROIs, and that it may be able to detect meaningful microstructural properties of such regions.

Standard neuroimaging techniques do not provide quantifiable data on brain microstructural architecture, however this study has demonstrated a reproducible and reliable method for obtaining whole brain maps with quantifiable estimates of brain microstructure. This measure is stable enough to be used in longitudinal studies lasting at least up to three months; and provides information on a voxel- or region-wise basis for analysis of subcortical structures, lesions, or developing brains (Dhollander et al., 2017; Mito et al., 2018; Mito et al., 2019; Bastiani et al., 2019). Related microstructural analysis of free water signal fractions has been performed in the context of Parkinson’s disease (Ofori et al., 2015; Burciu et al., 2016), Schizophrenia (Pasternak et al., 2012; Mandl et al., 2015), and concussion (Pasternak et al., 2014). 3-tissue CSD techniques may thus have the potential to be applied to a variety of these and other neurological conditions.

3-tissue CSD derived tissue fractions provide a flexible framework for analyzing diffusion images in ways not addressed in this paper. While we examined the reliability of WM/GM/CSF-like tissue signal fractions here, other researchers have used response functions representing different tissue compartments when contextually appropriate. Pietsch et al., (2019) applied two different WM response functions representing mature and immature WM in a developing adolescent cohort to observe WM maturation. Mito et al., (2019) proposed to apply a statistical framework of compositional data analysis to analyze the full 3-tissue composition of WM-, GM- and CSF-like signal fractions directly to study microstructure in white matter lesions, following the initial suggestion of moving towards such WM/GM/CSF-like diffusion signal fraction interpretation by Dhollander et al. (2017). In Aerts et al., (2019), this idea was furthermore used for the purpose of disentangling WM FODs representing infiltrated WM tracts in the presence of gliomas, so as to enable more reliable within-tumor tractography. Similar work has also recently been done by Chamberland et al., (2019), who illustrated the use of 3-tissue signal fractions in the presence of cerebral metastases, both to assess their microstructure as well as to enable tractography through nearby edematous regions.

The relatively recent use of CSD to describe the diffusion signal (Tournier et al., 2007) has led to some measure of controversy when compared to other established analysis techniques such as those based in multi-tensor models. One particular area of concern has been noted as the generation of ‘false-fibers’ on tracking algorithms due to spurious fODF peaks (Parker et al., 2013; Ning et al., 2015). Some studies using recent methodological improvements have suggested that the prevalence of false-fibers in CSD is oversold compared to other methods (Wilkins et al., 2015; Schilling et al., 2018). In this study 3-tissue CSD demonstrated good reliability across all compartments, suggesting that if false-positive fibers are present, they are not large enough component of the overall signal in each voxel to introduce significant amounts of variance in the corresponding signal fractions.

An additional benefit provided by 3-tissue CSD methods is in the potential for tissue type specific masking. The CSF-like compartment presented in this paper is calculated as CSF-like diffusion in tissue by relying on the other compartments to identify which voxels were ‘tissue’. Unlike a binary tissue segmentation based on T1 intensity, calculations of WM- and GM-like signal fraction compartments together were used to define voxels where ‘tissue’ composed a majority of signal from each voxel. This process relied exclusively on the single, native space diffusion image instead of reslicing and warping a separate structural image or atlas. Future studies might be able to take advantage of this approach by examining tissue compartment magnitudes inside voxels defined by the behavior of other tissue compartments. For example, tracking CSF-like (free water) tissue infiltration into voxels defined by the high proportion of WM-like tissue during aging or in certain pathological contexts.

## Conclusion

In this study, we performed a test-retest reliability and longer term stability analysis of the 3-tissue signal fractions as obtained from 3-tissue CSD techniques. We found that 3-tissue CSD techniques provide reliable and stable estimates of tissue microstructure composition, up to 3 months longitudinally in a control population. This forms an important basis for further investigations utilizing 3-tissue CSD techniques to track changes in microstructure across a variety of conditions.

## Acknowledgements

NITRC, NITRC-IR, and NITRC-CE have been funded in whole or in part with Federal funds from the Department of Health and Human Services, National Institute of Biomedical Imaging and Bioengineering, the National Institute of Neurological Disorders and Stroke, under the following NIH grants: 1R43NS074540, 2R44NS074540, and 1U24EB023398 and previously GSA Contract No. GS-00F-0034P, Order Number HHSN268200100090U

## References

1000 Functional Connectomes Project (http://www.nitrc.org/projects/fcon_1000/); Nathan S. Kline Institute for Psychiatric Research (NKI), enhanced NKI (eNKI) test–retest data.

Aerts, H., Dhollander, T., & Marinazzo, D. (2019). Evaluating the performance of 3-tissue constrained spherical deconvolution pipelines for within-tumor tractography. bioRxiv, 629873. doi:10.1101/629873

Albi, A., Pasternak, O., Minati, L., Marizzoni, M., Bartrés-Faz, D., Bargalló, N., … & Fiedler, U. (2017). Free water elimination improves test–retest reproducibility of diffusion tensor imaging indices in the brain: A longitudinal multisite study of healthy elderly subjects. Human brain mapping, 38(1), 12–26.

Alexander, A. L., Hasan, K. M., Lazar, M., Tsuruda, J. S., & Parker, D. L. (2001). Analysis of partial volume effects in diffusion-tensor MRI. Magnetic Resonance in Medicine, 45(5), 770–780.

Alexander, A. L., Lee, J. E., Lazar, M., & Field, A. S. (2007). Diffusion tensor imaging of the brain. Neurotherapeutics, 4(3), 316–329.

Andersson, J. L., & Sotiropoulos, S. N. (2016). An integrated approach to correction for off-resonance effects and subject movement in diffusion MR imaging. Neuroimage, 125, 1063–1078.

Andersson, J. L., Graham, M. S., Zsoldos, E., & Sotiropoulos, S. N. (2016). Incorporating outlier detection and replacement into a non-parametric framework for movement and distortion correction of diffusion MR images. NeuroImage, 141, 556–572.

Andersson, J. L., Skare, S., & Ashburner, J. (2003). How to correct susceptibility distortions in spin-echo echo-planar images: application to diffusion tensor imaging. Neuroimage, 20(2), 870–888.

Assaf, Y., & Pasternak, O. (2008). Diffusion tensor imaging (DTI)-based white matter mapping in brain research: a review. Journal of molecular neuroscience, 34(1), 51–61.

Avants, B. B., Tustison, N. J., Stauffer, M., Song, G., Wu, B., & Gee, J. C. (2014). The Insight ToolKit image registration framework. Frontiers in Neuroinformatics, 8, 44.

Avants, B. B., Tustison, N., & Song, G. (2009). Advanced normalization tools (ANTS). Insight j, 2, 1–35.

Basser, P. J., Mattiello, J., & LeBihan, D. (1994). MR diffusion tensor spectroscopy and imaging. Biophysical journal, 66(1), 259–267.

Bastiani, M., Andersson, J., Cordero-Grande, L., Murgasova, M., Hutter, J., Price, A. N., … & Victor, S. (2019). Automated processing pipeline for neonatal diffusion MRI in the developing Human Connectome Project. NeuroImage, 185(1), 750–763.

Biswal, B. B., Mennes, M., Zuo, X. N., Gohel, S., Kelly, C., Smith, S. M., … & Dogonowski, A. M. (2010). Toward discovery science of human brain function. Proceedings of the National Academy of Sciences, 107(10), 4734–4739.

Boullerne, A. I. (2016). The history of myelin. Experimental neurology, 283, 431–445.

Burciu, R. G., Ofori, E., Shukla, P., Pasternak, O., Chung, J. W., McFarland, N. R., … & Vaillancourt, D. E. (2016). Free-water and BOLD imaging changes in Parkinson’s disease patients chronically treated with a MAO-B inhibitor. Human brain mapping, 37(8), 2894–2903.

Chamberland, M., Iqbal, N. S., Rudrapatna, S. U., Parker, G., Tax, C. M., Staffurth, J., … & Jones, D. K. (2019). Characterising tissue heterogeneity in cerebral metastases using multi-shell multi-tissue constrained spherical deconvolution. Proceedings of the 27th International Society of Magnetic Resonance in Medicine, 27, 3613.

Chenevert, T. L., Brunberg, J. A., & Pipe, J. G. (1990). Anisotropic diffusion in human white matter: demonstration with MR techniques in vivo. Radiology, 177(2), 401–405.

Dell’Acqua, F., & Tournier, J. D. (2018). Modelling white matter with spherical deconvolution: How and why?. NMR in Biomedicine, e3945.

Dhollander, T., & Connelly, A. (2016). A novel iterative approach to reap the benefits of multi-tissue CSD from just single-shell (+ b= 0) diffusion MRI data. Proceedings of the 24th International Society of Magnetic Resonance in Medicine, 24, 3010.

Dhollander, T., Raffelt, D., & Connelly, A. (2016). Unsupervised 3-tissue response function estimation from single-shell or multi-shell diffusion MR data without a co-registered T1 image. In ISMRM Workshop on Breaking the Barriers of Diffusion MRI (Vol. 5).

Dhollander, T., Raffelt, D., & Connelly, A. (2017). Towards interpretation of 3-tissue constrained spherical deconvolution results in pathology. Proceedings of the 25th International Society of Magnetic Resonance in Medicine, 25, 1815.

Greenspan, H. (2008). Super-resolution in medical imaging. The Computer Journal, 52(1), 43–63.

Jenkinson, M., Beckmann, C. F., Behrens, T. E. J., Woolrich, M. W., & Smith, S. M. (2012). FSL. NeuroImage, 62(2), 782–790.

Jeurissen, B., Leemans, A., Tournier, J. D., Jones, D. K., & Sijbers, J. (2013). Investigating the prevalence of complex fiber configurations in white matter tissue with diffusion magnetic resonance imaging. Human brain mapping, 34(11), 2747–2766.

Jeurissen, B., Tournier, J. D., Dhollander, T., Connelly, A., & Sijbers, J. (2014). Multi-tissue constrained spherical deconvolution for improved analysis of multi-shell diffusion MRI data. NeuroImage, 103, 411–426.

Jiang, J., Zhao, Y. J., Hu, X. Y., Du, M. Y., Chen, Z. Q., Wu, M., … & Gong, Q. Y. (2017). Microstructural brain abnormalities in medication-free patients with major depressive disorder: a systematic review and meta-analysis of diffusion tensor imaging. Journal of psychiatry & neuroscience: JPN, 42(3), 150.

Johansen-Berg, H., & Behrens, T. E. (Eds.). (2013). Diffusion MRI: from quantitative measurement to in vivo neuroanatomy. Academic Press.

Jones, D. K., & Cercignani, M. (2010). Twenty-five pitfalls in the analysis of diffusion MRI data. NMR in Biomedicine, 23(7), 803–820.

Kellner, E., Dhital, B., Kiselev, V. G., & Reisert, M. (2016). Gibbs-ringing artifact removal based on local subvoxel-shifts. Magnetic resonance in medicine, 76(5), 1574–1581.

Kuklisova-Murgasova, M., Quaghebeur, G., Rutherford, M. A., Hajnal, J. V., & Schnabel, J. A. (2012). Reconstruction of fetal brain MRI with intensity matching and complete outlier removal. Medical image analysis, 16(8), 1550–1564.

Le Bihan, D., Mangin, J. F., Poupon, C., Clark, C. A., Pappata, S., Molko, N., & Chabriat, H. (2001). Diffusion tensor imaging: concepts and applications. Journal of Magnetic Resonance Imaging, 13(4), 534–546.

Mandl, R. C., Pasternak, O., Cahn, W., Kubicki, M., Kahn, R. S., Shenton, M. E., & Pol, H. E. H. (2015). Comparing free water imaging and magnetization transfer measurements in schizophrenia. Schizophrenia research, 161(1), 126–132.

Mito, R., Dhollander, T., Raffelt, D., Xia, Y., Salvado, O., Brodtmann, A., Rowe, C., Villemagne, V., & Connelly, A. (2018). Investigating microstructural heterogeneity of white matter hyperintensities in Alzheimer’s disease using single-shell 3-tissue constrained spherical deconvolution. Proceedings of the 26th International Society of Magnetic Resonance in Medicine, 26, 0135.

Mito, R., Dhollander, T., Xia, Y., Raffelt, D., Salvado, O., Churilov, L., … & Connelly, A. (2019). In vivo microstructural heterogeneity of white matter lesions in Alzheimer’s disease using tissue compositional analysis of diffusion MRI data. bioRxiv, 623124. doi:10.1101/623124

Nimsky, C., Ganslandt, O., Hastreiter, P., Wang, R., Benner, T., Sorensen, A. G., & Fahlbusch, R. (2005). Preoperative and intraoperative diffusion tensor imaging-based fiber tracking in glioma surgery. Neurosurgery, 56(1), 130–138.

Ning, L., Laun, F., Gur, Y., DiBella, E. V., Deslauriers-Gauthier, S., Megherbi, T., … & St-Jean, S. (2015). Sparse Reconstruction Challenge for diffusion MRI: Validation on a physical phantom to determine which acquisition scheme and analysis method to use?. Medical image analysis, 26(1), 316–331.

Nooner, K. B., Colcombe, S., Tobe, R., Mennes, M., Benedict, M., Moreno, A., … & Sikka, S. (2012). The NKI-Rockland sample: a model for accelerating the pace of discovery science in psychiatry. Frontiers in neuroscience, 6, 152.

Ofori, E., Dekosky, S. T., Febo, M., Colon-Perez, L., Chakrabarty, P., Durara, R., … & Vaillancourt, D. E. (2019). Free-water imaging of the hippocampus is a sensitive marker of Alzheimer’s disease. Neuroimage: Clinical, 101985.

Ofori, E., Pasternak, O., Planetta, P. J., Li, H., Burciu, R. G., Snyder, A. F., … & Vaillancourt, D. E. (2015). Longitudinal changes in free-water within the substantia nigra of Parkinson’s disease. Brain, 138(8), 2322–2331.

Oscar-Berman, M., & Marinković, K. (2007). Alcohol: effects on neurobehavioral functions and the brain. Neuropsychology review, 17(3), 239–257.

Parker, G. D., Marshall, D., Rosin, P. L., Drage, N., Richmond, S., & Jones, D. K. (2013). A pitfall in the reconstruction of fibre ODFs using spherical deconvolution of diffusion MRI data. Neuroimage, 65, 433–448.

Pasternak, O., Kelly, S., Sydnor, V. J., & Shenton, M. E. (2018). Advances in microstructural diffusion neuroimaging for psychiatric disorders. NeuroImage, 182(1), 259–282.

Pasternak, O., Koerte, I. K., Bouix, S., Fredman, E., Sasaki, T., Mayinger, M., … & Skopelja, E. N. (2014). Hockey Concussion Education Project, Part 2. Microstructural white matter alterations in acutely concussed ice hockey players: a longitudinal free-water MRI study. Journal of neurosurgery, 120(4), 873–881.

Pasternak, O., Sochen, N., Gur, Y., Intrator, N., & Assaf, Y. (2009). Free water elimination and mapping from diffusion MRI. Magnetic Resonance in Medicine, 62(3), 717–730.

Pasternak, O., Westin, C. F., Bouix, S., Seidman, L. J., Goldstein, J. M., Woo, T. U. W., … & Shenton, M. E. (2012). Excessive extracellular volume reveals a neurodegenerative pattern in schizophrenia onset. Journal of Neuroscience, 32(48), 17365–17372.

Pedraza, O., Bowers, D., & Gilmore, R. (2004). Asymmetry of the hippocampus and amygdala in MRI volumetric measurements of normal adults. Journal of the International Neuropsychological Society, 10(5), 664–678.

Pierpaoli, C., & Basser, P. J. (1996). Toward a quantitative assessment of diffusion anisotropy. Magnetic resonance in Medicine, 36(6), 893–906.

Pietsch, M., Christiaens, D., Hutter, J., Cordero-Grande, L., Price, A. N., Hughes, E., … & Tournier, J. D. (2019). A framework for multi-component analysis of diffusion MRI data over the neonatal period. NeuroImage, 186(1), 321–337.

Raffelt, D., Tournier, J. D., Fripp, J., Crozier, S., Connelly, A., & Salvado, O. (2011). Symmetric diffeomorphic registration of fibre orientation distributions. NeuroImage, 56(3), 1171–1180.

Raffelt, D., Tournier, J. D., Rose, S., Ridgway, G. R., Henderson, R., Crozier, S., … & Connelly, A. (2012). Apparent fibre density: a novel measure for the analysis of diffusion-weighted magnetic resonance images. Neuroimage, 59(4), 3976–3994.

Reynolds, B. B., Stanton, A. N., Soldozy, S., Goodkin, H. P., Wintermark, M., & Druzgal, T. J. (2017). Investigating the effects of subconcussion on functional connectivity using mass-univariate and multivariate approaches. Brain imaging and behavior, 1–14.

Rydhög, A. S., Szczepankiewicz, F., Wirestam, R., Ahlgren, A., Westin, C. F., Knutsson, L., & Pasternak, O. (2017). Separating blood and water: perfusion and free water elimination from diffusion MRI in the human brain. Neuroimage, 156, 423–434.

Schilling, K. G., Janve, V., Gao, Y., Stepniewska, I., Landman, B. A., & Anderson, A. W. (2018). Histological validation of diffusion MRI fiber orientation distributions and dispersion. NeuroImage, 165, 200–221.

Shattuck, D. W., Mirza, M., Adisetiyo, V., Hojatkashani, C., Salamon, G., Narr, K. L., … & Toga, A. W. (2008). Construction of a 3D probabilistic atlas of human cortical structures. Neuroimage, 39(3), 1064–1080.

Smith, S. M., Jenkinson, M., Woolrich, M. W., Beckmann, C. F., Behrens, T. E., Johansen-Berg, H., … & Niazy, R. K. (2004). Advances in functional and structural MR image analysis and implementation as FSL. Neuroimage, 23, S208–S219.

Tournier JD, Calamante F, Connelly A. (2007). Robust determination of the fibre orientation distribution in diffusion MRI: Non-negativity constrained super-resolved spherical deconvolution. NeuroImage, 35(4), 1459–1472.

Tournier, J. D., Mori, S., & Leemans, A. (2011). Diffusion tensor imaging and beyond. Magnetic resonance in medicine, 65(6), 1532–1556.

Tournier, J. D., Smith, R. E., Raffelt, D. A., Tabbara, R., Dhollander, T., Pietsch, M., … & Connelly, A. (2019). MRtrix3: A fast, flexible and open software framework for medical image processing and visualisation. bioRxiv, 551739. doi:10.1101/551739

Tuch, D. S., Reese, T. G., Wiegell, M. R., & Wedeen, V. J. (2003). Diffusion MRI of complex neural architecture. Neuron, 40(5), 885–895.

Veraart, J., Novikov, D. S., Christiaens, D., Ades-Aron, B., Sijbers, J., & Fieremans, E. (2016). Denoising of diffusion MRI using random matrix theory. NeuroImage, 142, 394–406.

Wiegell, M. R., Larsson, H. B., & Wedeen, V. J. (2000). Fiber crossing in human brain depicted with diffusion tensor MR imaging. Radiology, 217(3), 897–903.

Wilkins, B., Lee, N., Gajawelli, N., Law, M., & Leporé, N. (2015). Fiber estimation and tractography in diffusion MRI: development of simulated brain images and comparison of multi-fiber analysis methods at clinical b-values. Neuroimage, 109, 341–356.

Zhang, H., Schneider, T., Wheeler-Kingshott, C. A., & Alexander, D. C. (2012). NODDI: practical in vivo neurite orientation dispersion and density imaging of the human brain. Neuroimage, 61(4), 1000–1016.

